# Gene expression profiling in trigeminal ganglia from Cntnap2^-/-^ and Shank3b^-/-^ mouse models of autism spectrum disorder

**DOI:** 10.1101/2022.10.23.513403

**Authors:** Alessandra G. Ciancone-Chama, Yuri Bozzi, Luigi Balasco

## Abstract

Sensory difficulties represent a crucial issue in the life of autistic individuals. The diagnostic and statistical manual of mental disorders describes both hyper- and hypo-responsiveness to sensory stimulation as a criterion for the diagnosis autism spectrum disorders (ASD). Among the sensory domain affected in ASD, altered responses to tactile stimulation represent the most commonly reported sensory deficits. Although tactile abnormalities have been reported in monogenic cohorts of patients and genetic mouse models of ASD, the underlying mechanisms are still unknown. Traditionally, autism research has focused on the central nervous system as the target to infer the neurobiological bases of such tactile abnormalities. Nonetheless, the peripheral nervous system represents the initial site of processing of sensory information and a potential site of dysfunction in the sensory cascade. Here we investigated the gene expression deregulation in the trigeminal ganglion (which directly receives tactile information from whiskers) in two genetic models of syndromic autism (Shank3b and Cntnap2 mutant mice) at both adult and juvenile ages. We found several neuronal and non-neuronal markers involved in inhibitory, excitatory, neuroinflammatory and sensory neurotransmission to be differentially regulated within the trigeminal ganglia of both adult and juvenile Shank3b and Cntnap2 mutant mice. These results may help in entangling the multifaced complexity of sensory abnormalities in autism and open avenues for the development of peripherally targeted treatments for tactile sensory deficits exhibited in ASD.

## Introduction

Autism spectrum disorders (ASDs) form a multifaceted group of neurodevelopmental disorders with high heterogeneity of symptoms and severity. The diagnostic and statistical manual of mental disorders (DSM-V) describes autism as a pervasive neurological syndrome characterized by deficits in social interaction and repetitive/stereotyped behavior with several associated neurological symptoms. Despite being characterized by such a high degree of variability among individuals affected, it has been reported that 95% of autistic individuals show aberrant sensory experiences, suggesting sensory abnormalities as a typical feature in ASD. Indeed, sensory abnormalities are now described in the DSM-V as both hyper- and hypo-responsiveness to sensory stimulation demonstrating how sensory symptoms are fundamental for the description of the syndrome itself. Abnormal sensory reactivity represents a crucial issue in autism research since it likely contributes to other ASD symptoms such as anxiety, stereotyped behaviors, as well as cognitive and social dysfunctions (Ben-Sasson et al., 2007; Sinclair et al., 2017). Among the sensory domain affected in ASD, altered responses to tactile stimulation represent the most commonly reported sensory deficits in ASD (60,9%). Interestingly, hypo-responsiveness to tactile stimulation has been found to positively correlate with the severity of ASD core symptoms (Foss-Feig et al., 2012) and touch avoidance behavior in toddlers is predictive of ASD diagnosis later on in life (Mammen et al., 2015).

Although tactile abnormalities have been reported in monogenic cohorts of patients with ASD (Rett syndrome (Badr et al., 1987; Amir et al., 1999), fragile X syndrome (Rogers et al., 2003), Phelan-McDermid syndrome (Tavassoli et al. 2021), the underlying mechanisms are still under investigation. The first step in proper tactile perception starts in the periphery with the activation of the mechanosensory neurons which respond to innocuous tactile stimuli and mediate the perception of objects’ shape and texture, vibrations, and skin strokes. These neurons in mammals are pseudo-unipolar and have their cell bodies in the dorsal root ganglia (DRG) and trigeminal ganglia (TG), as part of the peripheral nervous system (PNS), where they mediate tactile sensitivity from the body and head region respectively. DRG and TG neurons collect the somatosensory information from the periphery and send projections to the central nervous system (CNS), within the spinal cord for DRG neurons, and the brainstem for TG neurons.

Thus, the DRG and TG are initial sites of processing of tactile stimuli then conveyed to CNS. Ideally, these two structures represent potential sites of dysfunction underlying impairments in somatosensory perception in ASD individuals. While defects in the DRG have been reported in mouse models of ASD (Orefice et al., 2016; Orefice et al., 2019), to the best of our knowledge there is no evidence of TG neuron dysfunctions in ASD.

Mouse lines harboring ASD-relevant mutations have been recently used to assess the neurobiological underpinnings of abnormal sensory responses in ASD within the central and the peripheral nervous system (He et al., 2017; Balasco et al., 2019, 2020; Chelini et al., 2019; Orefice, 2020; Pizzo et al., 2020). Given the relevance of the whisker system in mice, the assessment of TG neurons (which mediate tactile sensitivity from the whiskers) represents a promising tool to study somatosensory processing defects in ASD. Mice use their whiskers for a variety of behavior such as object exploration (Brecht, 2007) and conspecific interaction (Ahl, 1986), and abnormalities in sensory perception through whiskers profoundly impact mouse behavior (Arakawa and Erzurumlu, 2015; Erzurumlu and Gaspar, 2020).

In this work, we used *Shank3b* and *Cntnap2* mutant mice as models for the Phelan McDermid (PMS) and cortical dysplasia-focal epilepsy (CDFE) respectively, two syndromic forms of autism. PMS is caused by mutations to the SHANK3 gene which codes for the SH3 and multiple ankyrin repeat domain protein 3 (Monteiro & Feng, 2017). This protein belongs to the family of Shank proteins and therefore acts as a major scaffolding protein within the postsynaptic density of excitatory neurons (Jiang & Ehlers, 2013). As a syndrome, PMS is described by intellectual disability, speech and developmental delay, and importantly ASD-related behaviors such as problems in communication and social interaction (Phelan & McDermid, 2010), as well as sensory hypo reactivity to tactile stimulation (Balasco et al., 2020). CDFE is caused by a recessive nonsense mutation in the CNTNAP2 gene which codes for CASPR2. CASPR2 (Cntnap2) is part of the neurexin family of transmembrane proteins and is involved in neuron-glia interactions, potassium channel clustering on myelinated axons, dendritic arborization, and spine development (Poliak et al., 1999; Poliak et al., 2003; Anderson et al., 2012). CDFE syndrome is a rare disorder characterized by intellectual disability, ASD-like behaviors, language regression, and focal epileptic seizures from childhood Shank3b and Cntnap2 knockout mice display autistic-like characteristics as repetitive grooming and impaired social interaction among others thus are considered reliable models for ASDs (Peça et al., 2011; Peñagarikano et al., 2011; Vogt et al., 2018).

Given the fundamental role of early-life tactile experiences in shaping the acquisition of normal social behavior and communication skills in humans as well as in rodents, we hypothesized that peripheral tactile processing defects might contribute to ASD symptomatology. We recently reported aberrant somatosensory processing in the central nervous system of adult Shank3b and Cntnap2 mutant mice following whisker stimulation (Balasco et al., 2020; Balasco et al., 2021). Nonetheless, to infer a possible peripheral contribution to such defects, here we performed a gene expression profiling of TG in Cntnap2^-/-^ and Shank3b^-/-^ mice, in both adult and juvenile stages.

## Material and Methods

### Animals

All experiments were carried out following the Italian and European directives (DL 26/2014, EU 63/2010) and were approved by the Italian Ministry of Health and the University of Trento animal care committee. All surgical procedures were performed under anesthesia to minimize animal suffering. Animals were housed in a 12 h light/dark cycle with food and water available ad libitum. Shank3b and Cntnap2 mutant mice were crossed at least five times into a C57BL/6 background before mating. Heterozygous mating (Shank3b^+/-^ x Shank3b^+/-^ and Cntnap2+/- x Cntnap2+/-) was used to generate the wild-type (WT) and knockout (KO) homozygous littermates used in this study. PCR genotyping was performed following the Jackson Laboratory protocol (www.jax.org). 40 sex-matched adult (6 months old) littermates (10 Shank3b^+/+^ and 10 Shank3b^-/-^; 10 Cntnap2^+/+^ and 10 Cntnap2^-/-^) and 16 sex-matched juvenile (P30) littermates (3 Shank3b+/+ and 3 Shank3b^-/-^; 5 Cntnap2^+/+^ and 5 Cntnap2^-/-^) were used for qRT-PCR.

### Trigeminal Ganglia Dissection

Mice were anaesthetized and sacrificed by decapitation. Trigeminal ganglia (TG) were rapidly removed with surgical tweezers immediately after the skull’s opening and the brain’s removal. Ganglia were washed in cold PB buffer and stored at °80°C.

### Quantitative reverse transcription-polymerase chain reaction (RT-qPCR)

Total RNAs were extracted from the trigeminal ganglia of adult and young mutant (Shank3b^-/-^ and Cntnap2^-/-^) and control mice (Shank3b^+/+^ and Cntnap2^+/+^) with RNeasy Mini Kit (QUIAGEN) and retro-transcribed to cDNA with a SuperScript™ VILO™ cDNA Synthesis Kit (Invitrogen) according to the manufacturer’s protocol. qRT-PCR was performed in a CFX96TM Real-Time System (Bio-Rad, USA), using SYBR Green master mix (Bio-Rad). Primers (Sigma) were designed on different exons to avoid the amplification of genomic DNA (Table 1). The CFX3 Manager 3.0 (Bio-Rad) software was used to perform expression analyses (Sgadò et al., 2013). Mean cycle threshold (Ct) values from replicate experiments were calculated for each marker and β-actin (used as a standard for quantification), and then corrected for PCR efficiency and inter-run calibration. The expression level of each mRNA of interest was then normalized to that of β-actin for both genotypes. For RT-qPCR experiments, the expression levels of each marker (normalized to that of β-actin) were compared from triplicate experiments performed on RNA pools. Data can be found in supplementary tables (S2-5)

### Primers

Fourty eight genes were selected for testing via RT-qPCR on adult TG tissue, whereas a subset of 19 genes was selected for testing in juvenile TG tissue. Genes were selected to identify specific markers of sensory, inhibitory, and excitatory neurons, as well as neuroinflammation and neuroprotection. Primer specificity was verified using the In-Silico PCR (UCSC Genome Browser) and Primer Blast (NCBI) resources. Primer sequences are listed in Supplementary Table 1 (S1).

### Gene expression Analysis

All raw Ct values resulting from qRT-PCR experiments were normalised and calculated into Fold change values via the Livak method. Fold change (FC) represents the expression ratio of KO groups (Shank3b^-/-^ or Cntnap2^-/-^) relative to their respective WT controls (Shank3b^+/+^ or Cntnap2^+/+^). Therefore, fold changes lower than 1 represent a downregulation in the expression of a target gene while those greater than 1 report an upregulation of the target gene relative to the control sample. Statistical analysis was performed by unpaired t-test using the GraphPad Prism 8 software with a significance level set at p<0.05.

## Results

Figure 1 describes the experimental workflow used to profile gene expression in trigeminal ganglia from Shank3b and Cntnap2 mutant mice. A total of 48 genes were tested in the trigeminal ganglia of WT and KO *Shank3b* and *Cntnap2* adult mice. A subset of 19 genes were selected post-hoc and chosen for testing in trigeminal ganglia from juvenile mutant mice and controls (P30 WT and KO).

**Fig. 1).**
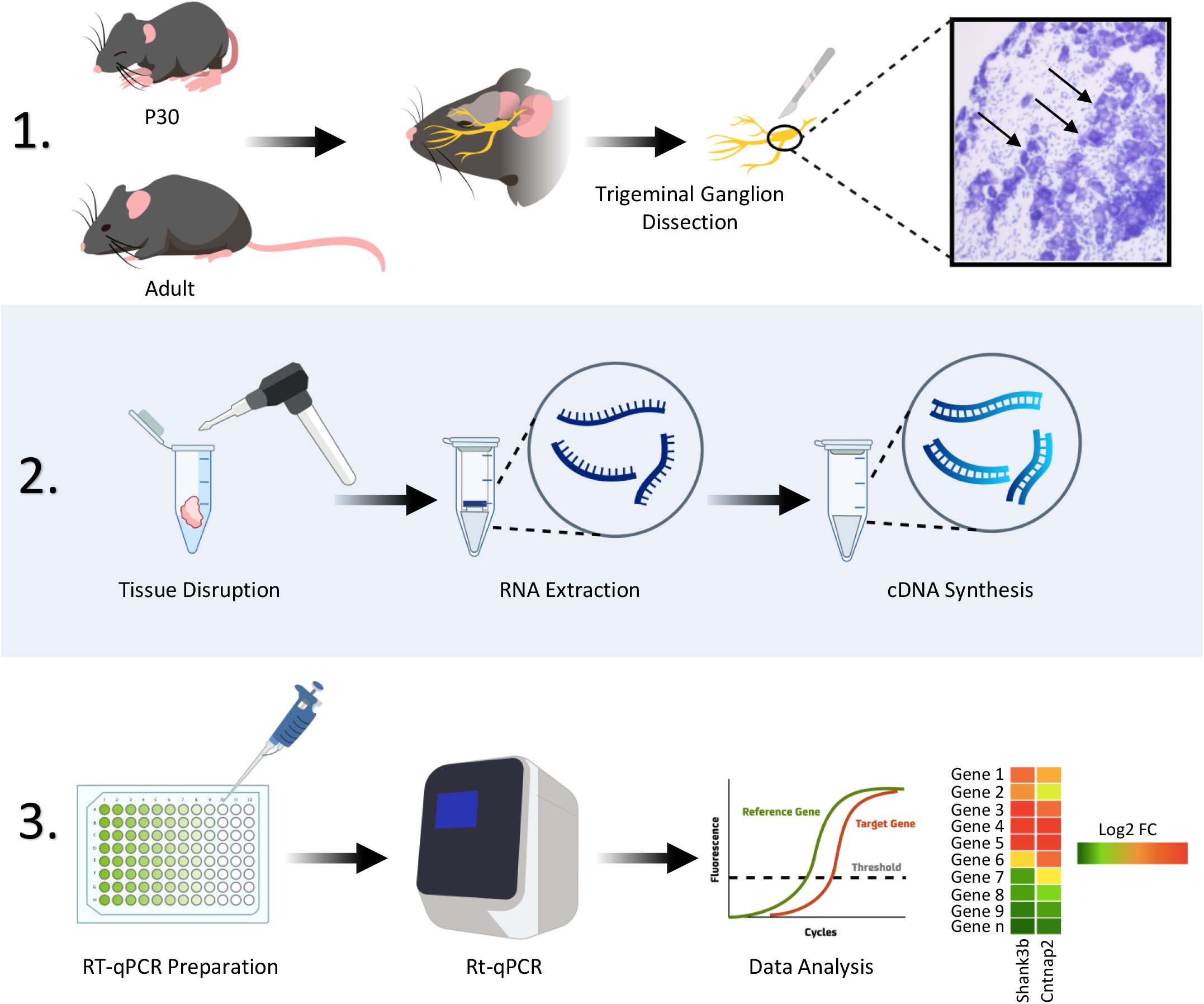
Schematic of the experimental workflow. Adult and P30 mice of both Shank3b and Cntnap2 mice were sacrificed and trigeminal ganglia (TG) collected. A Nissl staining of the TG is presented. Arrows indicates pseudo-unipolar neurons (in light blue). TG is then mechanically disrupted and the RNA extracted and converted in cDNA. RT-qPCR is performed and run on a Real-time PCR thermal cycler. Data are extracted and analyzed as described.

### Adult mice

Numerous markers for sensory neurons were significantly upregulated in the trigeminal ganglia of either Shank3b^-/-^ or Cntnap2^-/-^ adult mice (Fig 2A). Cck (FC=2.164, p=0.0001), Sst (FC=1.387, p=0.0333), P2X7 (FC=1.201, p=0.0378), and Calca (FC=1.222, p=0.0019) show significant increases in expression values in Cntnap2^-/-^ trigeminal ganglia, similarly as Tacr1 (FC= 1.435, p=0.0006) and Trpa1 (FC= 1.154, p=0.0185) in Shank3b^-/-^ samples. As in Cntnap2 KO TG, Calca mRNA was also upregulated in Shank3b^-/-^ TG (FC=1.114, p=0.0144).

**Fig. 2).**
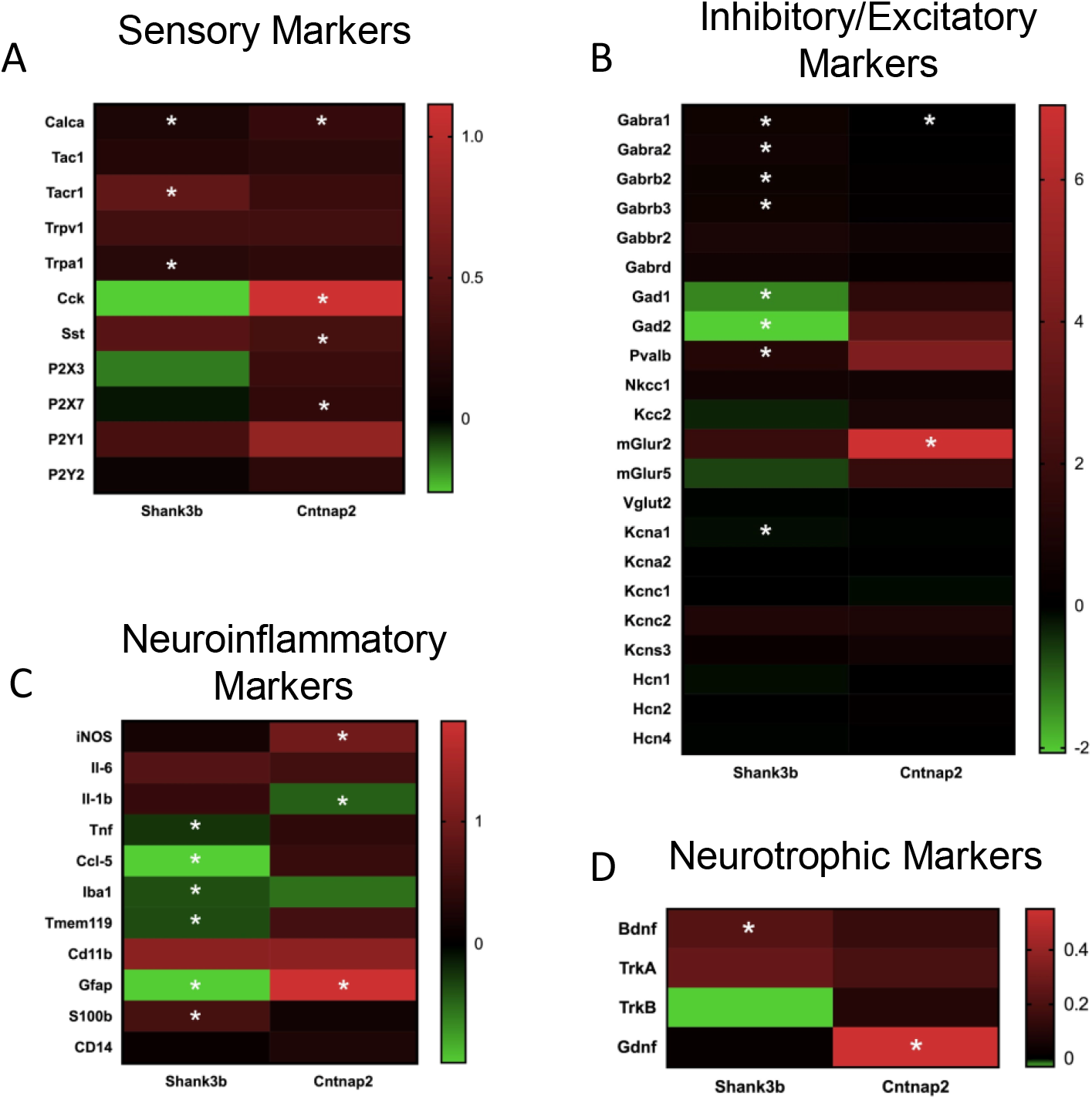
Heatmap of Log2 fold change values for adult Shank3b and Cntnap2 TG. Values above 0 indicate an upregulation of the respective gene in KO mice relative to WT controls (in red). Conversely values below 0 indicate a downregulation of a gene in KO animals relative to WT controls (in green). Genes annotated with an asterisk (*) show a significant difference in expression values (p < 0.05).

Shank3b^-/-^ and Cntnap2^-/-^ adult mice also showed significant alteration in the expression of inhibitory markers (Fig 2B) such as a downregulation in Gabra1 (for SHK FC=1.435, p<0.0001; for CNT FC= 1.072, p=0.0038) in the trigeminal ganglia. There was a significant decrease in the genetic expression of Gad2 (FC = 0.2379, p< 0.0001) in the TG of Shank3b^-/-^ adults. On the other hand, genes such as Pvalb (FC=2.313, p< 0.0001), Gabra2 (FC=1.549, p< 0.0001), Gabrb2 (FC=1.399, p=0.0021), and Gabrb3 (FC=1.450, p=0.0434) were upregulated in the TG of these same mice. Markers for excitatory neuronal subtypes were also significantly differentially expressed in the trigeminal ganglia of KO adult animals (Fig 2B). In Shank3b KO mice, Kcna1 was downregulated (FC= 0.8985, p= 0.0447). Additionally, an upregulation of mGluR2 (FC=132.0, p= 0.0103) was found in Cntnap2 KO animals.

Several markers for neuroinflammation (Fig 2C) were downregulated in the adult Shank3b^-/-^ TG, namely CCL-5 (FC=0.5115, p=0.0012), Tnf (FC=0.8403, p=0.0226), Iba1 (FC=0.7728, p=0.0476), and Tmem-119 (FC=0.7799, p=0.0103). qRT-PCR showed an increased expression of S100B mRNA in Shank3b KO TG (FC=1.519, p=0.0321). In *Cntnap2^-/-^* mice there was also an upregulation of iNOS expression (FC=1.864, p=0.0003). Conversely, Il-1b expression showed to be significantly decreased in Cntnap2^-/-^ samples (FC=0.7344, p=0.0023).

Finally, a significant increase in the gene expression was found among the neurotrophic markers tested in adult samples (Fig 2D), with TrkA (FC=1.205, p= 0.0332) showing an upregulation in Shank3b^-/-^ samples and Gdnf (FC=1.464, p=0.0404) in Cntnap2^-/-^ samples.

### Juvenile mice

qRT-PCR experiments on juvenile mice showed that only Shank3b^-/-^ TGs display a significant difference in the expression of sensory markers, namely a decrease in P2X3 mRNA expression (FC= 0.6916, p=0.0031) and an increased expression of Cck (FC= 4.087, p=0.0220).

Within the inhibitory markers tested, Gabrd was upregulated in Shank3b^-/-^ trigeminal ganglia (FC=1.241, p 0.0016), while mGlur5 is upregulated in Shank3b^-/-^ (FC=2.277, p= 0.0494) among the excitatory markers tested.

Of the neuroinflammatory markers tested in P30 mice, Il-1b was the only one resulting in a significant difference in fold change showing an upregulation (FC= 1.817, p= 0.0107) in Cntnap2^-/-^ juvenile mice relative to controls.

Lastly, in the category of neurotrophic markers, both TrkA and TrkB were downregulated (FC= 0.7446, p= 0.0472 for TrkA; FC= 0.8958, p=0.0175 for TrkB), while Gdnf was upregulated (FC= 2.086, p=0.0053) in the Shank3b^-/-^ TG.

#### Gad1 and Gfap

The expression profile of Gad1 and GFAP was of particular interest as these genes showed differential expression in both adult and juvenile Shank3b^-/-^ and Cntnap2^-/-^ mice. At P30, Gad1 expression in both Shank3b^-/-^ and Cntnap2^-/-^ TG was significantly higher relative to WT controls (FC=2.193, p=0.0103 for Shank3b^-/-^; FC=1.750, p=0.0397 for Cntnap2^-/-^) (Fig 4A,B). In adulthood, a significant downregulation of these gene was found only in the case of Shank3b^-/-^ TG (FC=0.3933, p<0.0001) (Fig 4C, D).

**Fig. 3).**
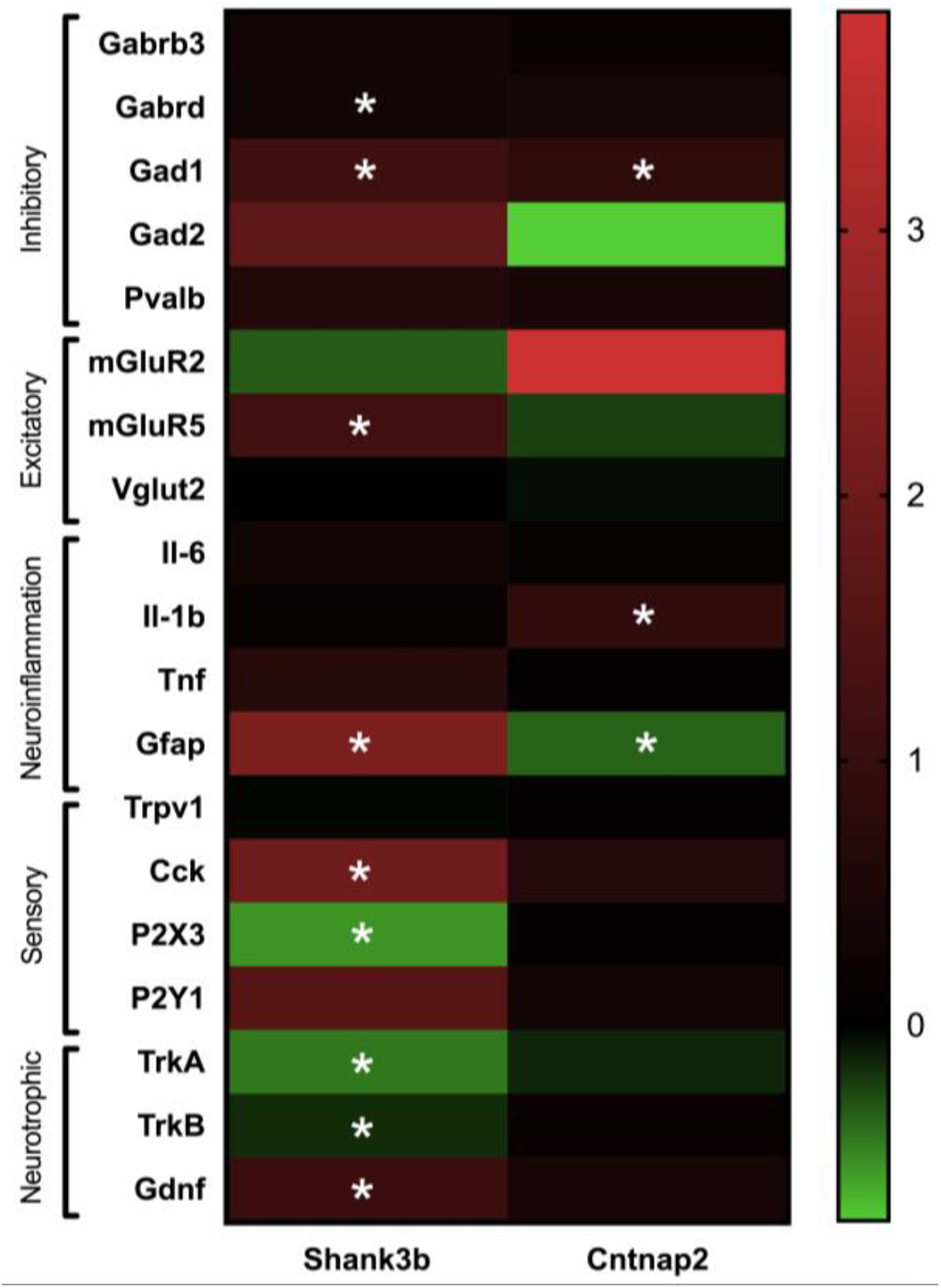
Heatmap of Log2 fold change values for juvenile (P30) Shank3b and Cntnap2 TG. Values above 0 indicate an upregulation of the respective gene in KO mice relative to WT controls (in red). Conversely values below 0 indicate a downregulation of a gene in KO animals relative to WT controls (in green). Genes annotated with an asterisk (*) show a significant difference in expression values (p < 0.05).

**Fig. 4).**
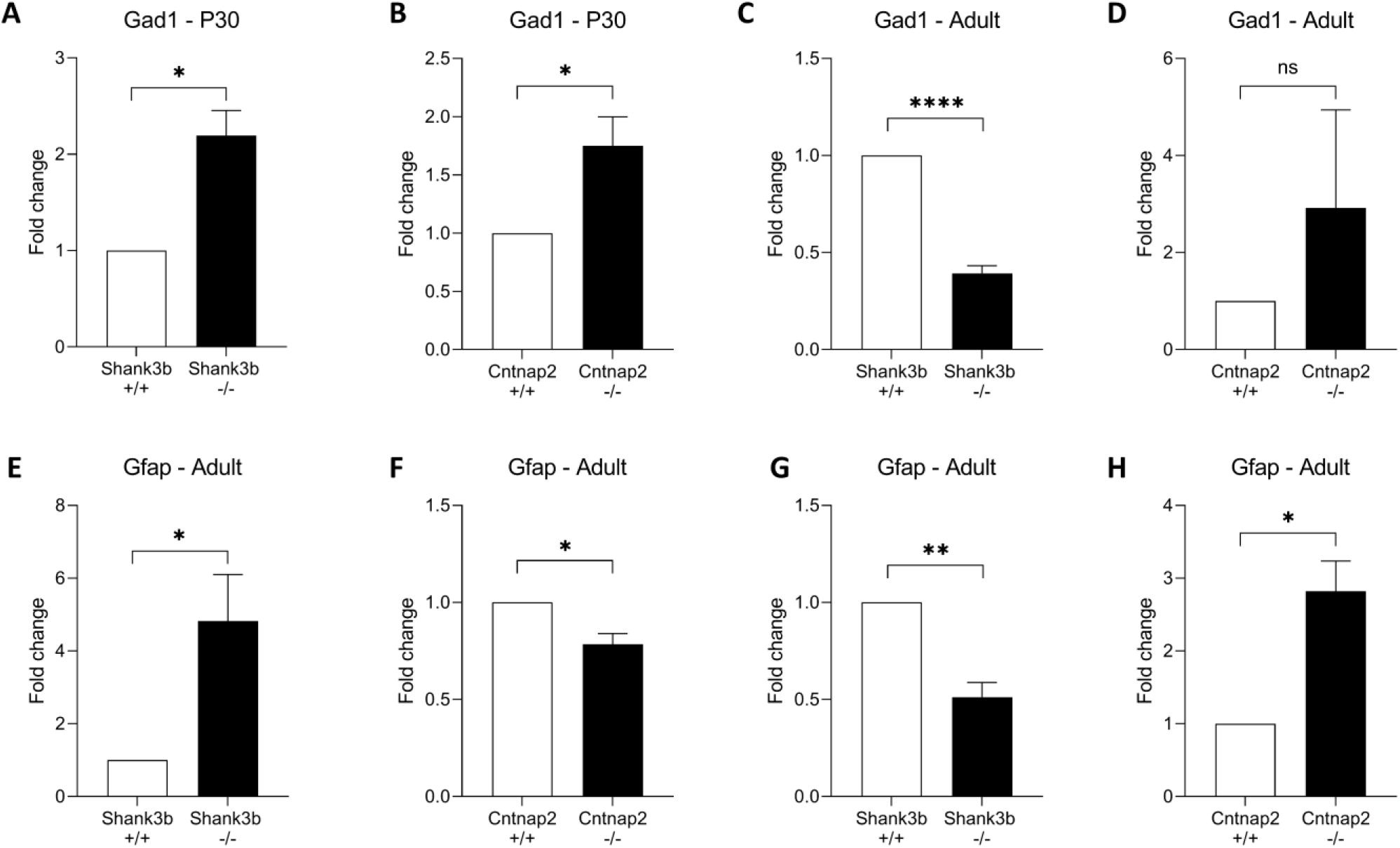
Fold change values for Gad1 and Gfap in Shank3b and Cntnap2 adult and juvenile (P30) TG.

Gfap expression showed opposed trends throughout development in the two mouse strains. Gfap mRNA was upregulated at P30 (FC=4.823, p=0.0403) and downregulated in adulthood (FC=0.5118, p=0.0030) in Shank3b^-/-^ mice (Fig 4E, F). In contrast, juvenile Cntnap2^-/-^ mice showed Gfap downregulation (FC=0.7848, p=0.0173) followed by its upregulation in adulthood (FC=2.821, p=0.0117) (Fig 4G, H).

## Discussion

The TG is among the first stations in tactile sensory processing in the mouse somatosensory system. Altered expression of key gene markers within this crucial area may hence affect the ability for proper sensory experiences and thus outcome in behavioral abnormalities. The present investigation sought to test several neuronal and non-neuronal markers involved in cellular mechanisms (inhibition, excitation, neuroinflammation) within the trigeminal ganglia of both adult and juvenile Shank3b and Cntnap2 mouse models of ASD. Untangling the role of such markers based solely on their gene expression throughout development carries an inherent level of difficulty as several cellular processes may interfere with and influence the resulting behavioral phenotypes. Nonetheless, the present work is the first to describe gene expression alteration in trigeminal ganglia of genetic mouse models of ASD and give possible support for the involvement of particular markers within the TG in ASD.

The present study revealed significant alterations in markers for sensory neurons in the TG of Shank3b^-/-^ and Cntnap2^-/-^ mice, even at juvenile ages suggesting possible abnormalities in sensory neuron structure and functioning early on. Specifically, we showed an increased mRNA expression in several sensory markers involved in nociception, such as Trap1, Calca, and Tacr1 in Shank3b^-/-^ and Calca and Sst in Cntnap2^-/-^ TG. Furthermore, P2X7, whose inhibition was found to ameliorate dendritic spine pathology and social deficits in Rett syndrome mice models (Garré et al., 2020) is upregulated in Cntnap2^-/-^ adults. Such findings converge with previous studies reporting instances of increased pain sensitivity in ASD patients and animal models (Zhang et al., 2021; Failla et al., 2020). Other researchers however find a hyposensitivity in response to stimuli among ASD individuals and models (Dhamne et al., 2017; Allely, 2013). Altered expression of sensory markers P2X3 and Cck in juvenile Shank3b^-/-^ mice (downregulated and upregulated respectively) attest to early changes in sensory marker expression which could possibly contribute to the sensory dysfunction. The purinergic P2X3 receptor (associated with nociception and hypersensitivity) is upregulated by inflammation, oxidative stress, pain, (Zhang et al., 2014; Wu et al., 2004) and epilepsy (Zhou et al., 2016), all of which have been documented in relation to ASD. A deregulated expression of these markers could directly affect the initial stages of sensory processing thereafter leading to hyper- or hyposensitivity typically seen among ASD mouse models.

Dysregulation in GABAergic marker expression and interneuron functioning is commonly seen among humans with ASD as well as within animal models of this disorder (Zhao et al., 2022; Cellot & Cherubini, 2014; Vogt et al., 2018). In line with this, the present study found that inhibitory markers, the majority of which GABAergic, were among the most altered in Shank3b^-/-^ and Cntnap2^-/-^ TG. This general finding supports the excitatory/inhibitory imbalance theory, which postulates that the prevailing force of excitation over inhibition contributes to core ASD symptomology (Rubenstein & Merzenich, 2003). In this effect, previous studies note the reduction of GABA receptor units such as GABAA receptors in various cortical areas using diverse models of ASD (Zhao et al., 2022; Cellot & Cherubini, 2014; Fatemi et al., 2014; Orefice et al., 2019). These results are in stark contrast to the present results which highlight an increase of GABAergic markers such as Gabra1, Gabra2, Gabrb2, and Gabrb3 in Shank3b^-/-^ and Cntnap2^-/-^ trigeminal ganglia. It should be noted that a similar upregulation in mRNA expression has been shown in the cerebellum and parietal cortex of individuals with ASD (Fatemi et al., 2014). Upregulation found in GABAergic markers in the trigeminal ganglia can be consequently interpreted as underlining increased inhibition and thus giving way to hyporeactive/hyposensitive behaviors in adult Shank3b^-/-^ and Cntnap2^-/-^ mice (Balasco et al., 2020; Balasco et al., 2021). In juvenile mice, on the other hand, an upregulation was found among Shank3b^-/-^ TG for Gabrd, a gene whose dysfunction promotes psychomotor delay and epilepsy (Windpassinger et al., 2002). Interestingly and in agreement with our qRT-PCR result, gain-of-function variants of Gabrd have been associated with neurodevelopmental disorders (Ahring et al., 2022).

Our qRT-PCR results also unveiled an upregulation of the metabotropic glutamate receptor 5 (mGluR5) in the trigeminal ganglia of Shank3b^-/-^ juvenile mice. Upregulation of this gene was likewise found in the vermis of young ASD individuals post-mortem (Fatemi et al., 2011). The role of mGluR5 in ASD is supported by studies that associate its increased activity with the manifestation of symptoms in Fragile X Syndrome (Yan et al., 2005). Likewise, blocking mGluR5 activity via antagonists results in the reversal of core ASD symptoms (Mehta et al., 2011). Another noteworthy result is the downregulation of Kcna1 in Shank3b^-/-^ adult TG which encodes for the Kv1.1 voltage-gated potassium channel subunits that regulate axonal excitability and whose loss of function missense mutations are often associated with ataxia and epilepsy (Thouta et al., 2021). Kcna^-/-^ mice have been shown to demonstrate normal sociability and reduced repetitive behaviors as compared to WT counterparts (Indumathy et al., 2021). These results, suggesting an increased excitability in both Shank3b and Cntnap2 models, seem to conflict with our findings on inhibitory markers. It is imperative to underline, however, that contrasting evidence on the role of excitatory markers in ASD is available (Oka & Takashima, 1999; Lohith et al., 2013; Chana et al., 2015; Cai et al., 2019) and that the present results are of smaller magnitude than that of other marker types. Hence, further investigation is crucial for untangling the influence of excitatory gene expression in ASD.

Neuroinflammation has been proposed to contribute the development of ASDs by prompting neuronal dysfunction that characterize these conditions (Eissa et al., 2020). Several lines of research document the elevated expression of inflammatory markers such as pro-inflammatory cytokines in the brain and blood of ASD subjects (Varga et al., 2005; Zhao et al., 2021; Li et al., 2009; Rodriguez & Kern, 2011; Molloy et al., 2006). Our experiments showed an upregulation of inflammatory markers such as S100b, iNOS, and Il-1b in the TG of Shank3b^-/-^ and Cntnap2^-/-^ mice. Other inflammatory markers (Tnf, Iba1, Ccl-5, and Tmem-119) were instead downregulated in the TG of Shank3b^-/-^ and Cntnap2^-/-^ mice. While no evidence of immune dysfunction in the trigeminal ganglia from ASD individuals or animal models have been reported so far, our data could signal a state of immunosuppression. As ASD individuals often exhibit a dysregulated immune system vulnerable to viral insults (Sabourin et al., 2019; Hughes et al., 2018) a decreased production of pro-inflammatory proteins could contribute to immunodeficiencies. Likewise, there is the possibility that persistent inflammation early in development, instigated by overexpression of pro-inflammatory cytokines and chemokines, may promote a compensatory anti-inflammatory overresponse later on. Such speculations require further investigation in order to understand the role of neuroinflammation as a product and/or instigator underlying neuronal dysfunction in the context of the peripheral nervous system in ASD.

Finally, neurotrophic factors are crucial for neural development throughout our lifespan but especially during critical periods. Consequently, these growth factors have been implicated in the origins of neurodevelopmental disorders such as ASD and ADHD (Nickl-Jockschat & Michel, 2011). Of these neurotrophic factors, Bdnf is widely investigated in relation to ASD with numerous studies documenting either an increase or decrease (Nickl-Jockschat & Michel, 2011; Skogstrand et al., 2019) in the brains of ASD individuals. Apart from differing by brain regions (Louhivuori et al., 2011), alterations in Bdnf levels also depend on the severity of behavioral deficits (Kasarpalkar et al., 2014). While Bdnf mRNA was not found to be altered in either model in the present study, TrkB was significantly downregulated in the TG of juvenile Shank3b^-/-^mice. Bdnf exerts its effect through coupling with its receptor, TrkB, and this signaling pathway is believed to be defective (hyperactivated) in ASD (Subramanian et al., 2015). In fact, reports of the administration of TrkB agonist or partial agonist rescuing ASD-related symptoms provide support for its involvement in ASD neuropathology (Lee & Han, 2019; Tsai, 2006). Ngf (nerve growth factor) is another neurotrophic factor crucial to typical development which has been found to be altered in ASD (Mostafa et al., 2021; Dinçel et al., 2013). Again, while Ngf itself was not altered in either model, our results do show a downregulation of its receptor TrkA in Shank3b^-/-^ juvenile TG. TrkA mediates Ngf effects such as cell differentiation, proliferation, and programmed death (Martin-Zanca et al., 1986). Thus, the downregulation of these neurotrophic receptors may compromise the function of crucial factors in early development. The glial-derived nerve growth factor Gdnf was likewise found to be upregulated in Shank3b^-/-^ juvenile and Cntnap2^-/-^ adult TG. Gdnf, whose upregulation is associated with ADHD (Galvez-Contreras et al., 2017), supports cell survival (specifically of dopaminergic cells) (Lin et al., 1993) and thus may contribute to the dysregulation of excitatory synaptic activity in the TG.

Among the markers tested in the present study, the expression profiles of Gad1 and Gfap mRNA are particularly relevant. While Gad1 is downregulated in Shank3b^-/-^ adults, its expression is upregulated in KO juveniles of both lines. Gad1 encodes for the GAD67 enzyme crucial to GABA synthesis, particularly during early neurodevelopment (Feldblum et al., 1993), and its dysregulation may be involved in creating an excitatory/inhibitory imbalance (Rubenstein & Merzenich, 2003). Lower levels of both Gad1 mRNA and protein have been found in the postmortem brains of autistic adults (Fatemi et al., 2002; Chao et al., 2010; Yip et al., 2007; Zhubi et al., 2017) as well as in animal models (Peñagarikano et al., 2011) in several areas throughout the cerebral cortex. On the other hand, increased Gad1 expression has been documented in the cerebellum of postmortem autistic brains (Yip et al., 2008) and the prefrontal cortex of ASD animal models (Hou et al., 2018; El Idrissi et al., 2005). Gad1 overexpression correlates with increased GABA synthesis and therefore promotes inhibition (Dicken et al., 2015). Chemogenetic depolarization of GABAergic DRG reduces peripherally-induced nociception, while decreasing inhibition by introducing GABA receptor antagonists on sensory ganglia trigger nociception (Du et al., 2017). Moreover, recent work on cortical networks (Haroush & Marom, 2019) suggests that inhibition may reduce discrimination between stimuli. Thus, upregulation of Gad1 in the TG of juvenile Shank3b^-/-^ and Cntnap2^-/-^ mice may lead to an increase in inhibition from sensory neurons upstream to higher-level sensory cortex leading to altered texture discrimination, as observed in these mice (Balasco et al., 2021, 2022).

The present study also found Gfap mRNA upregulation in the juvenile Shank3b^-/-^ and in the adult Cntnap2^-/-^ TG. Conversely, Gfap mRNA was downregulated in the TG of Shank3b^-/-^ adults and Cntnap2^-/-^ juveniles. The Gfap gene codes for GFAP expressed in glial cells, in the TG specifically in satellite glial cells (Stephenson & Byers, 1995). GFAP expression has been reported to be deregulated in the brain of autistic subjects (Laurence & Fatemi, 2005; Edmonson et al., 2014Crawford et al., 2015). Therefore, alterations in Gfap expression observed in juvenile and adult Shank3b^-/-^ and Cntnap2^-/-^ may reflect disturbances in astrocyte function that could compromise typical neural development and transmission as seen in ASD (Petrelli et al., 2016).

In this study, we uncovered for the first time significant differences in gene expression of important markers of neuronal and glial function in the TG of Shank3b^-/-^ and Cntnap2^-/-^ mice at young and adult age. A true understanding of the functional significance of these alterations in gene expression found in the TG of Shank3b^-/-^ and Cntnap2^-/-^ will only be achieved prior further investigations. Such results open avenues for the development of peripherally targeted treatments for tactile sensory deficits observed in ASD.

## Supporting information

Ciancone et al. 2022_Suppl. 1

Ciancone et al. 2022_Suppl. 2

Ciancone et al. 2022_Suppl. 3

Ciancone et al. 2022_Suppl. 4

Ciancone et al. 2022_Suppl. 5

## Conflict of Interest

The authors declare that the research was conducted in the absence of any commercial or financial relationships that could be construed as a potential conflict of interest.

## Author Contributions

AGCC performed experiments, analysed data and drafted the manuscript; YB provided funding and edited the manuscript; LB designed and supervised the study, analyzed data, and wrote the manuscript.

## Funding

This work was supported by the Strategic Project TRAIN—Trentino Autism Initiative (https://projects.unitn.it/train/index.html) from the University of Trento (grant 2018–2021 to YB). LB is a recipient of a PhD fellowship from the University of Trento and Fondazione CARITRO (Trento, Italy)

## Acknowledgments

We thank the technical and administrative staff of CIMeC for the excellent assistance and Simona Correale for her precious help with figure preparation.

## Supplementary Material

Supplementary Table 1. Primers used for quantitative RT-PCR experiments.

Supplementary Table 2. RT-qPCR results from Shank3b adult TG

Supplementary Table 3. RT-qPCR results from Cntnap2 adult TG

Supplementary Table 4. RT-qPCR results from Shank3b juvenile TG (P30)

Supplementary Table 5. RT-qPCR results from Cntnap2 juvenile TG (P30)

## Notes

### Competing Interest Statement

The authors have declared no competing interest.

