## Supplementary material for "Gene expression profiling in trigeminal ganglia from Cntnap2^-/-^ and Shank3b^-/-^ mouse models of autism spectrum disorder": Ciancone et al. 2022_Suppl. 1

| Gene Symbol | Gene name | Forward Sequence 5’->3’ | Reverse Sequence 5’->3’ |
|---|---|---|---|
| **Beta Actin** | Beta Actin | AATCGTGCGTGACATCAAAG | AAGGAAGGCTGGAAAAGAGC |
| **Cntnap2** | Contactin Associated Protein 2 | GAGGGGAGAAAAAGCAAAGCAG | AACTGCTGTACCTTCCTTGGG |
| **Shank3b** | SH3 And Multiple Ankyrin Repeat Domains 3 | TTACACCCACACCTGCCTTC | CACCATCCTCCTCGGGTTTC |
| **Gad1** | Glutamate Decarboxylase 1 | AGATAGCCCTGAGCGACGAG | ATGGCCGATGATTCTGGTTC |
| **Gabra2** | GABA A Receptor Subunit Alpha 2 | AGATTCAAAGCCACTGGAGG | CCAGCACCAACCTGACTG |
| **Gabrb3** | GABA A Receptor Subunit Beta 3 | GAGGTCTTCACAAGCTCAAAATC | AGGCAGGGTAATATTTCACTCAG |
| **Gabra1** | GABA A Receptor Subunit Alpha 1 | CTCTCCCACACTTTTCTCCC | CCGACAGTGTGCTCAGAATG |
| **Gabrb2** | GABA A Receptor Subunit Beta 2 | TCAGAGGATGACTTTGCTA | GCACACAATAATGTTTACTAT |
| **Gabbr2** | GABA B Receptor Subunit Beta 2 | TCAGAGGATGACTTTGCTA | GCACACAATAATGTTTACTAT |
| **Gabrd** | GABA A Receptor Subunit Delta | ATGCATTTGCCCACTTCAA | ATGGGTTTGAGTCTGGAACG |
| **Gad2** | Glutamate Decarboxylase 2 | CATTCCTGTCCTTGCCTCTC | GTGCATCCTTTGTCCATGT |
| **Pvalb** | GABAergic Interneuron Subpopulation | TGTCGATGACAGACGTGCTC | TTCTTCAACCCCAATCTTGC |
| **Nkcc1** | SLC12A1 - Solute Carrier Family 12 Member 1 | CCTCTCACGAACCCATTGG | GCTGGGATAGGTCTCTCTGT |
| **Kcc2** | SLC12A1 - Solute Carrier Family 12 Member 4 | AGATCGAGAGCAACGACGAGAGG | GGTGGCGATCGAAGAAGAAT |
| **Calca** | Alpha CGRP Peptide | CCTGCAACACTGCCACCTGCG | GAAGGCTTCAGAGCCCACATTG |
| **Tac1** | Tachykinin Precursor 1 (Substance P) | ATGGCCAGATCTCTCACAAAAG | AAGATGAATAGATAGTGCGTTCAGG |
| **Tacr1** | Tachykinin Precursor 1 Receptor (Substance P-R) | GCTCTGTGCATGGGTCTCTT | AGGAAGGATGGCTCCAGGAT |
| **Trpv1** | Transient Receptor Potential Vanilloid 1 | CAAACTCCACCCCACACTGA | AGGCCAAGACCCCAATCTTC |
| **Trpa1** | Transient Receptor Potential Cation Channel | AGGTGATTTTTAAAACATTGCTGAG | CTCGATAATTGATGTCTCCTAGCAT |
| **Cck** | Cholecystokinin | AGCGGCGTATGTCTGTGCGT | CACTGCGCCGGCCAAAATCC |
| **Sst** | Somatostatin | CCCCAGACTCCGTCAGTTTCT | TCTCTGTCTGGTTGGGCTCG |
| **P2X3** | Purinergic Receptor P2X 3 | CAGGGCACTTCTGTCTTTGTC | AGCGGTACTTCTCCTCATTCTC |
| **P2X7** | Purinergic Receptor P2X 7 | CGAGTTGGTGCCAGTGTGGA | CCTGCTGTTGGTGGCCTCTT |
| **P2Y1** | Purinergic Receptor P2Y 1 | CCTGCGAAGTTATTTCATCTA | GTTGAGACTTGCTAGACCTCT |
| **P2Y2** | Purinergic Receptor P2Y 2 | GCAGCATCCTCTTCCTCACCT | CATGTTGATGGCGTTGAGGGT |
| **Cckar** | Cholecystokinin Receptor A | GCTGCATAGCGTCACTTGG | GATGGAGTTAGACTGCAACC |
| **Cckbr** | Cholecystokinin Receptor B | CCAAGCTGCTGGCTAAGAAG | CTTAGCCTGGACAGAGAAGC |
| **Cacna1c** | Calcium Voltage-Gated Channel Subunit Alpha1 C | CGTTCTCATCCTGCTCAACACC | GAGCTTCAGGATCATCTCCACTG |
| **Ramp1** | Receptor (calcitonin) Activity Modifying Protein 1 | GACGCTATGGTGTGACT | AGTGCAGTCATGAGCAG |
| **Calcrl** | Calcitonin Receptor-like | TGCTGGAATGACGTTGCAGC | GCCTTCACAGAGCATCCAGA |
| **Il-6** | Interleukin-6 | GCCTTCTTGGGACTGATGCT | GACAGGTCTGTTGGGAGTGG |
| **Il-1b** | Interleukin-1b | ACGGACCCCAAAAGATGAAG | TTCTCCACAGCCACAATGAG |

| | | | |
|---|---|---|---|
| **iNOS** | Nitric Oxide Synthase 2, citokine - Inducible | CAGCTGGGCTGTACAAACCTT | CATTGGAAGTGAAGCGTTTCG |
| **Tnf** | Tumor Necrosis Factor | CAAAATTCGAGTGACAAGCC | TGTCTTTGAGATCCATGCCG |
| **Ccl-5** | C-C Motif Chemokine Ligand 5 | AGAATACATCAACTATTTGGAGA | CCTTGCATCTGAAATTTTAATGA |
| **Iba1** | AIF1 - Allograft Inflammatory Factor 1 | GTCCTTGAAGCGAATGCTGG | CATTCTCAAGATGGCAGATC |
| **Tmem119** | Transmembrane Protein 119 | GTGTCTAACAGGCCCCAGAA | AGCCACGTGGTATCAAGGAG |
| **Cd11b** | IGAM - Integrin Subunit Alpha M | CCTTGTTCTCTTTGATGCAG | GTGATGACAACTAGGATCTT |
| **Gfap** | Glial Fibrillary Acidic Protein | TCCTGGAACAGCAAAACAAG | CAGCCTCAGGTTGGTTTCAT |
| **S100b** | S100 Calcium Binding Protein B | TGCCCTCATTGATGTCTTCCA | GAGAGAGCTCGTTGTTGATAAGCT |
| **CD14** | CD14 Molecule | GGCTTGTTGCTGTTGCTTC | CAGGGCTCCGAATAGAATCC |
| **CD68** | CD68 Molecule | AGGGTGGAAGAAAGGTAAAGC | AGAGCAGGTCAAGGTGAACAG |
| **mGlurR5** | Metabotropic Glutamate Receptor 5 | ATCTGCCTGGGTTACTTGTG | GCAATACGGTTGGTCTTCG |
| **mGluR2** | GRM2 - Glutamate Metabotropic Receptor 2 | AGGCCATGCTTTTTGCACTG | GAAGGCCTCAATGCCTGTCT |
| **Vglut2** | Vescicular Glutamatergic Transporter 2 | TGCTACCTCACAGGAGAATGGA | GCGCACCTTCTTGCACAAAT |
| **Kcna1** | Voltage-Gated Potassium Channel Subunit Kv1.1 | TTACCCTGGGCACGGAGATA | ACACCCTTACCAAGCGGATG |
| **Kcna2** | Voltage-Gated Potassium Channel Subunit Kv1.2 | CATCTGCAAGGGCAACGTCAC | CCTTTGGAAGGAAGGAGGCA |
| **Kcns3** | Voltage-Gated Potassium Channel Subunit Kv9.3 | CCCTGGACAAGATGAGGAAC | TTGATGCCCCAGTACTCGAT |
| **Hcn2** | Hyperpolarization Activated Cyclic Nucleotide Gated Potassium Channel 2 | ATCGCATAGGCAAGAAGAACTC | CAATCTCCTGGATGATGGCATT |
| **Hcn1** | Hyperpolarization Activated Cyclic Nucleotide Gated Potassium Channel 1 | CTCAGTCTCTTGCGGTTATTACG | TGGCGAGGTCATAGGTCAT |
| **Kcnc1** | Voltage-Gated Potassium Channel Subunit Kv3.1 | GTGCCGACGAGTTCTTCTTC | GTCATCTCCAGCTCGTCCTC |
| **Vglut1** | Vescicular Glutamatergic Transporter 1 | CCCCCAAATCCTTGCACTTT | AACAAATGGCCACTGAGAAACC |
| **Knc2c** | Voltage-Gated Potassium Channel Subunit Kv3.2 | AGATCGAGAGCAACGAGAGG | GGTGGCGATCGAAGAAGAAT |
| **Hcn4** | Hyperpolarization Activated Cyclic Nucleotide Gated Potassium Channel 4 | GCATGATGCTTCTGCTGTGT | GCTTCCCCCAGGAGTTATTC |
| **Bdnf** | Brain Derived Neurotrophic Factor | AGGCCAACTGAAGCAGTATTTC | CCGAACATACGATTGGGTAGTT |
| **TrkA** | Neurotrophic Receptor Tyrosine Kinase 1 | GAAGAATGTGACGTGCTGGG | GAAGGAGACGCTGACTTGGA |
| **TrkB** | Neurotrophic Receptor Tyrosine Kinase 2 | AAGGACTTTCATCGGGAAGCTG | TCGCCCTCCACACAGACAC |
| **Gdnf** | Glial Cell Derived Neurotrophic Factor | GCCACCATTAAAAGACTGAAAAGG | GCCTGCCGATTCCTCTCTCT |

Supplementary Table 1. Primers used for quantitative RT-PCR experiments.
