## Supplementary material for "Gene expression profiling in trigeminal ganglia from Cntnap2^-/-^ and Shank3b^-/-^ mouse models of autism spectrum disorder": Ciancone et al. 2022_Suppl. 2

| Target gene | Fold change 2^-(ΔΔCt) Mean | p-value | p-value summary | Difference between means (B - A) ± SEM |
|---|---|---|---|---|
| **Calca** | 1.114 | 0.0144 | * | 0.1138 ± 0.02749 |
| **Tac1** | 1.144 | 0.0719 | ns | 0.1437 ± 0.05912 |
| **Tacr1** | 1.435 | 0.0006 | *** | 0.4354 ± 0.04519 |
| **Trpv1** | 1.275 | 0.466 | ns | 0.2747 ± 0.3530 |
| **Trpa1** | 1.154 | 0.0185 | * | 0.1540 ± 0.04013 |
| **Cck** | 0.8354 | 0.4178 | ns | -0.1646 ± 0.1824 |
| **Sst** | 1.296 | 0.2897 | ns | 0.2965 ± 0.2553 |
| **P2X3** | 0.8981 | 0.4514 | ns | -0.1019 ± 0.1223 |
| **P2X7** | 0.9804 | 0.9033 | ns | -0.01957 ± 0.1545 |
| **P2Y1** | 1.297 | 0.7041 | ns | 0.2969 ± 0.7451 |
| **P2Y2** | 1.052 | 0.8368 | ns | 0.05242 ± 0.2437 |
| **iNOS** | 1.138 | 0.7351 | ns | 0.1381 ± 0.3806 |
| **Il-6** | 1.656 | 0.3884 | ns | 0.6563 ± 0.7059 |
| **Il-1b** | 1.387 | 0.3261 | ns | 0.3866 ± 0.3615 |
| **Tnf** | 0.8403 | 0.0226 | * | -0.1598 ± 0.05243 |
| **Ccl-5** | 0.5115 | 0.0012 | ** | -0.4885 ± 0.05970 |
| **Iba1** | 0.7728 | 0.0476 | * | -0.2272 ± 0.08041 |
| **Tmem119** | 0.7799 | 0.0103 | * | -0.2201 ± 0.04820 |
| **Cd11b** | 2.334 | 0.4689 | ns | 1.334 ± 1.726 |
| **Gfap** | 0.5118 | 0.0030 | ** | -0.4882 ± 0.07603 |
| **S100b** | 1.519 | 0.0321 | * | 0.5191 ± 0.1609 |
| **CD14** | 1.068 | 0.8333 | ns | 0.06757 ± 0.3009 |
| **Gabra1** | 1.435 | <0.0001 | **** | 0.4345 ± 0.007424 |
| **Gabra2** | 1.549 | <0.0001 | **** | 0.5491 ± 0.02172 |
| **Gabrb2** | 1.399 | 0.0021 | ** | 0.3986 ± 0.05638 |
| **Gabrb3** | 1.45 | 0.0434 | * | 0.4503 ± 0.1544 |
| **Gabbr2** | 1.879 | 0.278 | ns | 0.8785 ± 0.7366 |
| **Gabrd** | 1.534 | 0.4518 | ns | 0.5342 ± 0.6415 |
| **Gad1** | 0.3933 | <0.0001 | **** | -0.6067 ± 0.03844 |
| **Gad2** | 0.2379 | <0.0001 | **** | -0.7621 ± 0.02352 |
| **Pvalb** | 2.313 | <0.0001 | **** | 1.313 ± 0.04096 |
| **Nkcc1** | 1.59 | 0.6401 | ns | 0.5900 ± 1.168 |
| **Kcc2** | 0.7812 | 0.7046 | ns | -0.2188 ± 0.5371 |
| **mGluR2** | 3.692 | 0.44 | ns | 2.692 ± 3.143 |
| **mGluR5** | 0.6241 | 0.0986 | ns | -0.3759 ± 0.1752 |
| **Vglut2** | 0.9549 | 0.0777 | ns | -0.04513 ± 0.01913 |
| **Kcna1** | 0.8985 | 0.0447 | * | -0.1015 ± 0.03515 |
| **Kcna2** | 1.022 | 0.6314 | ns | 0.02240 ± 0.04320 |
| **Kcnc1** | 1.004 | 0.9555 | ns | 0.004100 ± 0.06898 |
| **Kcnc2** | 2.096 | 0.1758 | ns | 1.096 ± 0.7520 |
| **Kcns3** | 1.336 | 0.6924 | ns | 0.3364 ± 0.7907 |
| **Hcn1** | 0.905 | 0.2502 | ns | -0.09497 ± 0.07067 |
| **Hcn2** | 1.033 | 0.8234 | ns | 0.03337 ± 0.1401 |
| **Hcn4** | 0.9572 | 0.6982 | ns | -0.04283 ± 0.1028 |
| **Bdnf** | 1.167 | 0.454 | ns | 0.1674 ± 0.2021 |
| **TrkA** | 1.205 | 0.0332 | * | 0.2054 ± 0.06436 |
| **TrkB** | 0.98 | 0.9117 | ns | -0.01997 ± 0.1692 |
| **Gdnf** | 1.012 | 0.9407 | ns | 0.01220 ± 0.1540 |

Supplementary Table 2. RT-qPCR results from Shank3b adults
