## Supplementary material for "Gene expression profiling in trigeminal ganglia from Cntnap2^-/-^ and Shank3b^-/-^ mouse models of autism spectrum disorder": Ciancone et al. 2022_Suppl. 3

| Target gene | Fold change 2^-(ΔΔCt) Mean | p-value | p-value summary | Difference between means (B - A) ± SEM |
|---|---|---|---|---|
| **Calca** | 1.222 | 0.0019 | ** | 0.2217 ± 0.04202 |
| **Tac1** | 1.175 | 0.4072 | ns | 0.1752 ± 0.1893 |
| **Tacr1** | 1.235 | 0.3087 | ns | 0.2352 ± 0.2018 |
| **Trpv1** | 1.278 | 0.4706 | ns | 0.2777 ± 0.3608 |
| **Trpa1** | 1.187 | 0.6365 | ns | 0.1871 ± 0.3761 |
| **Cck** | 2.164 | 0.0001 | *** | 1.164 ± 0.1706 |
| **Sst** | 1.387 | 0.0333 | * | 0.3865 ± 0.1213 |
| **P2X3** | 1.239 | 0.2042 | ns | 0.2394 ± 0.1579 |
| **P2X7** | 1.201 | 0.0378 | * | 0.2009 ± 0.06575 |
| **P2Y1** | 1.731 | 0.4071 | ns | 0.7311 ± 0.8202 |
| **P2Y2** | 1.176 | 0.5836 | ns | 0.1761 ± 0.3040 |
| **iNOS** | 1.864 | 0.0003 | *** | 0.8638 ± 0.07301 |
| **Il-6** | 1.491 | 0.4748 | ns | 0.4911 ± 0.6442 |
| **Il-1b** | 0.7344 | 0.0023 | ** | -0.2656 ± 0.03850 |
| **Tnf** | 1.334 | 0.3068 | ns | 0.3337 ± 0.2988 |
| **Ccl-5** | 1.4 | 0.3793 | ns | 0.4005 ± 0.4056 |
| **Iba1** | 0.6926 | 0.3195 | ns | -0.3074 ± 0.2707 |
| **Tmem119** | 1.497 | 0.0812 | ns | 0.4973 ± 0.2144 |
| **Cd11b** | 2.36 | 0.4609 | ns | 1.360 ± 1.726 |
| **Gfap** | 2.821 | 0.0117 | * | 1.821 ± 0.4140 |
| **S100b** | 1.112 | 0.7462 | ns | 0.1120 ± 0.3305 |
| **CD14** | 1.2 | 0.6769 | ns | 0.2005 ± 0.4580 |
| **Gabra1** | 1.072 | 0.0038 | ** | 0.07190 ± 0.01191 |
| **Gabra2** | 1.018 | 0.5518 | ns | 0.01783 ± 0.02748 |
| **Gabrb2** | 1.083 | 0.1299 | ns | 0.08263 ± 0.04345 |
| **Gabrb3** | 1.088 | 0.1462 | ns | 0.08763 ± 0.04867 |
| **Gabbr2** | 1.482 | 0.446 | ns | 0.4823 ± 0.5915 |
| **Gabrd** | 1.186 | 0.7205 | ns | 0.1863 ± 0.4851 |
| **Gad1** | 2.921 | 0.3954 | ns | 1.921 ± 2.020 |
| **Gad2** | 7.626 | 0.4165 | ns | 6.626 ± 7.319 |
| **Pvalb** | 19.96 | 0.3539 | ns | 18.96 ± 18.09 |
| **Nkcc1** | 1.524 | 0.646 | ns | 0.5237 ± 1.056 |
| **Kcc2** | 1.839 | 0.5377 | ns | 0.8392 ± 1.247 |
| **mGluR2** | 132 | 0.0103 | * | 131.0 ± 39.25 |
| **mGluR5** | 3.391 | 0.4415 | ns | 2.391 ± 2.801 |
| **Vglut2** | 1.02 | 0.7337 | ns | 0.01963 ± 0.05380 |
| **Kcna1** | 0.9735 | 0.4848 | ns | -0.02650 ± 0.03447 |
| **Kcna2** | 0.9928 | 0.8795 | ns | -0.007200 ± 0.04458 |
| **Kcnc1** | 0.9137 | 0.3746 | ns | -0.08633 ± 0.08647 |
| **Kcnc2** | 1.975 | 0.185 | ns | 0.9745 ± 0.6846 |
| **Kcns3** | 1.566 | 0.5381 | ns | 0.5664 ± 0.8421 |
| **Hcn1** | 1.02 | 0.8807 | ns | 0.01977 ± 0.1236 |
| **Hcn2** | 1.12 | 0.2014 | ns | 0.1204 ± 0.07882 |
| **Hcn4** | 1.036 | 0.7134 | ns | 0.03627 ± 0.09194 |
| **Bdnf** | 1.108 | 0.6029 | ns | 0.1080 ± 0.1916 |
| **TrkA** | 1.143 | 0.1953 | ns | 0.1430 ± 0.09203 |
| **TrkB** | 1.071 | 0.6811 | ns | 0.07143 ± 0.1615 |
| **Gdnf** | 1.464 | 0.0404 | * | 0.4636 ± 0.1552 |

Supplementary Table 3. RT-qPCR results from Cntnap2 adults
