## Supplementary material for "Gene expression profiling in trigeminal ganglia from Cntnap2^-/-^ and Shank3b^-/-^ mouse models of autism spectrum disorder": Ciancone et al. 2022_Suppl. 4

| Target gene | Fold change 2^-(ΔΔCt) Mean | p-value | p-value summary | Difference between means (B - A) ± SEM |
|---|---|---|---|---|
| **Trpv1** | 0.983 | 0.9004 | ns | -0.01700 ± 0.1275 |
| **Cck** | 4.087 | 0.022 | * | 3.087 ± 0.8480 |
| **P2X3** | 0.6916 | 0.0031 | ** | -0.3084 ± 0.04852 |
| **P2Y1** | 2.93 | 0.2738 | ns | 1.930 ± 1.523 |
| **Il-6** | 1.218 | 0.4243 | ns | 0.2182 ± 0.2455 |
| **Il-1b** | 1.109 | 0.675 | ns | 0.1086 ± 0.2405 |
| **Tnf** | 1.616 | 0.1344 | ns | 0.6163 ± 0.3291 |
| **Gfap** | 4.823 | 0.0403 | * | 3.823 ± 1.278 |
| **Gabrb3** | 1.276 | 0.5762 | ns | 0.2764 ± 0.4548 |
| **Gabrd** | 1.241 | 0.0016 | ** | 0.2405 ± 0.03153 |
| **Gad1** | 2.193 | 0.0103 | * | 1.193 ± 0.2614 |
| **Gad2** | 3.256 | 0.1993 | ns | 2.256 ± 1.468 |
| **Pvalb** | 1.539 | 0.6521 | ns | 0.5387 ± 1.108 |
| **mGluR2** | 0.7954 | 0.3631 | ns | -0.2046 ± 0.1995 |
| **mGluR5** | 2.277 | 0.0494 | * | 1.277 ± 0.4580 |
| **Vglut2** | 1.000 | 0.9974 | ns | 0.0003333 ± 0.09696 |
| **TrkA** | 0.7446 | 0.0472 | * | -0.2554 ± 0.09014 |
| **TrkB** | 0.8958 | 0.0175 | * | -0.1042 ± 0.02671 |
| **Gdnf** | 2.086 | 0.0053 | ** | 1.086 ± 0.1970 |

Supplementary Table 4. RT-qPCR results from Shank3b P30
