## Supplementary material for "Gene expression profiling in trigeminal ganglia from Cntnap2^-/-^ and Shank3b^-/-^ mouse models of autism spectrum disorder": Ciancone et al. 2022_Suppl. 5

| Target gene | Fold change 2^-(ΔΔCt) Mean | p-value | p-value summary | Difference between means (B - A) ± SEM |
|---|---|---|---|---|
| **Trpv1** | 1.033 | 0.7802 | ns | 0.03253 ± 0.1090 |
| **Cck** | 1.557 | 0.1954 | ns | 0.5565 ± 0.3584 |
| **P2X3** | 1.049 | 0.7009 | ns | 0.04930 ± 0.1194 |
| **P2Y1** | 1.245 | 0.7153 | ns | 0.2447 ± 0.6249 |
| **Il-6** | 1.068 | 0.8011 | ns | 0.06833 ± 0.2538 |
| **Il-1b** | 1.817 | 0.0107 | * | 0.8169 ± 0.1809 |
| **Tnf** | 1.028 | 0.9424 | ns | 0.02767 ± 0.3598 |
| **Gfap** | 0.7848 | 0.0173 | * | -0.2152 ± 0.05495 |
| **Gabrb3** | 1.152 | 0.7759 | ns | 0.1516 ± 0.4978 |
| **Gabrd** | 1.318 | 0.1135 | ns | 0.3179 ± 0.1574 |
| **Gad1** | 1.750,0 | 0.0397 | * | 0.7497 ± 0.2494 |
| **Gad2** | 0.6003 | 0.2817 | ns | -0.3997 ± 0.3215 |
| **Pvalb** | 1.352 | 0.7154 | ns | 0.3518 ± 0.8984 |
| **mGluR2** | 14.18 | 0.1373 | ns | 13.18 ± 7.686 |
| **mGluR5** | 0.8529 | 0.5532 | ns | -0.1471 ± 0.2276 |
| **Vglut2** | 0.9664 | 0.6684 | ns | -0.03363 ± 0.07286 |
| **TrkA** | 0.9177 | 0.5226 | ns | -0.08230 ± 0.1176 |
| **TrkB** | 1.085 | 0.4703 | ns | 0.08483 ± 0.1065 |
| **Gdnf** | 1.342 | 0.1677 | ns | 0.3419 ± 0.2032 |

Supplementary Table 5. RT-qPCR results from Cntnap2 P30
